# Cyclic uremia without kidney injury drives cardiovascular disease with rapid onset of immunosenescence and heart failure

**DOI:** 10.64898/2026.09.29.755324

**Authors:** Gideon JL Schaefer, Patrick Droste, Carla Schikarski, Xinhui Li, Marius Kohl, Niklas Lutterbach, Henriette de Loor, Lars Koch, Sylvia Menzel, Ling Zhang, Qingqing Long, Kristian Vogt, Anne-Sophie Andries, Anne Babler, David Schumacher, KM Schneider, Andreas Hoeft, Peter Boor, Bjorn Meijers, Rebekka K Schneider, Carolin V. Schneider, Axel Honné, Rafael Kramann, Konrad Hoeft

## Abstract

End-stage kidney disease (ESKD) is a major driver of cardiovascular disease (CVD) and requires dialysis treatment leading to a unique metabolic phenotype with cyclic transitions between accumulation and clearance of uremic toxins ^1–3^. Although ESKD patients display the highest cardiovascular mortality among CKD patients, the effect of cyclic uremia on cardiovascular disease remains subject to discussion ^4–6^. Existing mouse models rely on induction of kidney injury and do not resolve the interconnected pathophysiological effects of metabolic, hypertensive and endocrine renal failure ^7–9^. They are therefore unable to distinguish uremia-driven effects from well-established drivers of cardiovascular disease such as hypertension. Here, we established a simple, reproducible, and sex-inclusive mouse model of uremia in absence of kidney injury or hypertension leveraging a bistable vesico-peritoneal shunt (VPS). Strikingly, we identify cyclic uremia as an independent driver of ESKD-induced CVD, that induces heart failure with preserved ejection fraction (HFpEF), vascular inflammation and immunosenescence. As such, the VPS represents the first preclinical model that enables investigation of cyclic uremia as a driver of cardiovascular disease.

## Main

CKD-associated mortality culminates in ESKD patients, who harbor a 28-fold increase in cardiovascular mortality compared to the general population (**Fig. 1a, Fig S1a**). In contrast to stable CKD, intermittent dialysis in ESKD patients leads to a unique metabolic profile shifting between fluid overload and accumulation of uremic toxins (uremia) due to kidney failure and normalization of the latter during dialysis (**Fig. 1b**). These fluctuations are not captured by existing CKD mouse models that either mimic stable (e.g. 5/6 nephrectomy) or progressive (e.g. adenine diet) CKD, or AKI-to-CKD transition (e.g. bilateral ischemia reperfusion injury) (**Fig S1b**) ^9^. Existing rodent CKD-models are based on kidney injury, leading to disruption of the endocrine and exocrine kidney function and uremia remains a bystander beneath CKD-induced hypertension and endocrine dysregulation in those models ^7–9^. As a consequence, the effect of uremia as an independent driver of CVD can not be resolved in-vivo. Notably, a recent benchmark highlighted that these models fail to capture key features of ESKD-CVD, such as cardiac fibrosis ^9^.

**Figure 1:**
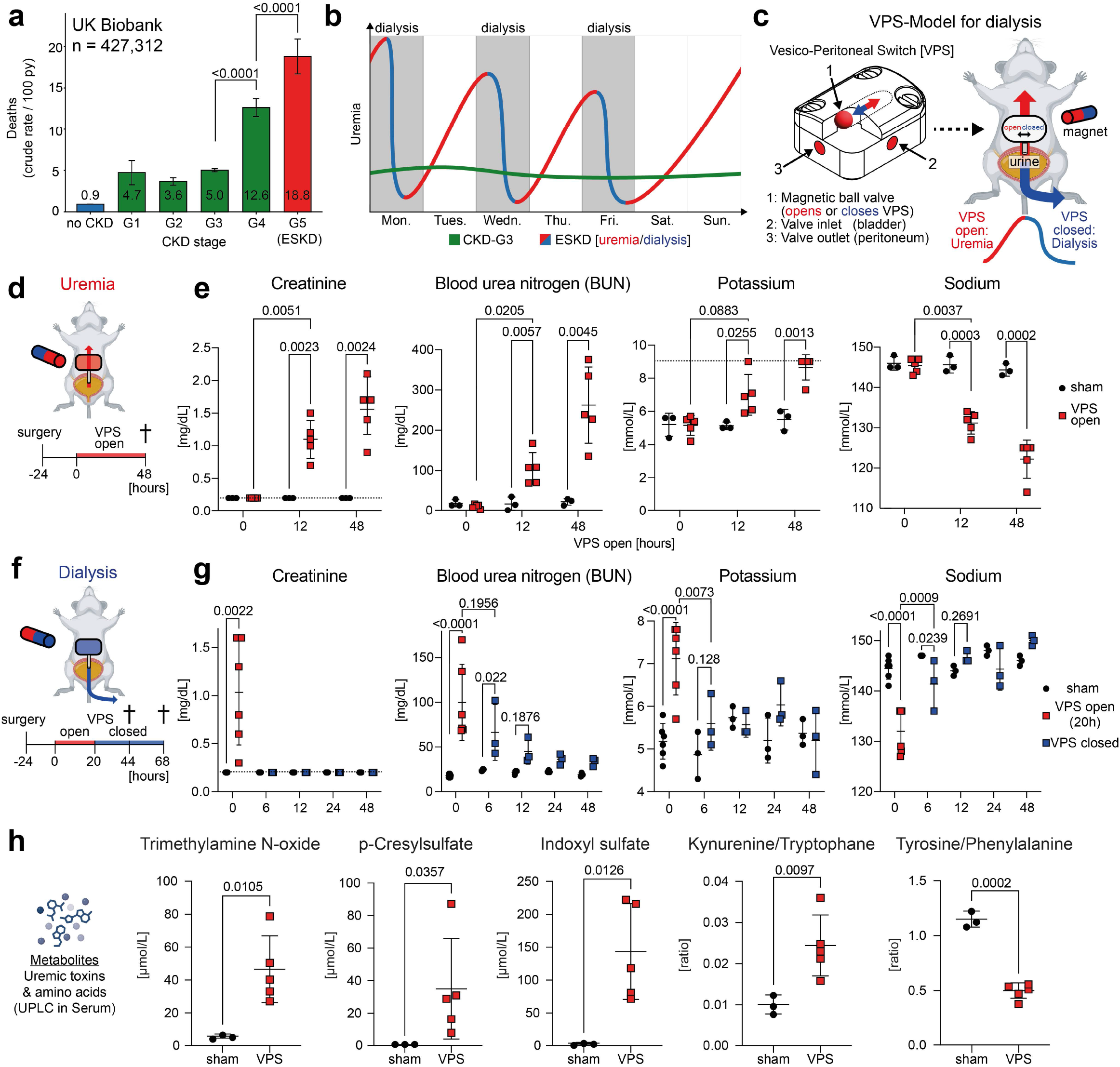
VPS enables induction and resolution of uremia. **a)** Bar chart indicating the death rate per 100 patient years in 427,312 patients of the UK biobank stratified by CKD stage. For CKD staging, the first ICD code assigned for CKD (N18.1–N18.5) was used. Individuals who had undergone a kidney transplant (Z94.0 or T86.1) were excluded from the analysis. **b)** Schematic diagram showing the temporal progression of dialysis-specific cyclic uremia in patients with CKD Stage 3 and CKD Stage 5 requiring dialysis (ESKD). **c)** Schematic of the vesicoperitoneal shunt (VPS) model. Implantation of a VPS with a bistable magnetically actuated valve enables discrete control of uremia [VPS open] and dialysis [VPS closed]. The valve consists of a ferromagnetic ball that is held in one of two states (open or close) by two magnets. The two VPS states (open/close) can be changed transdermally using an external magnet. When the VPS is open, urine flows into the peritoneum, which induces uremia through peritoneal resorption. Closing the VPS interrupts retrograde urine flow, causing it to be excreted transurethrally, corresponding to dialysis with clearance of uremia. **d)** Schematic diagram of uremia induction. Twenty-four hours after switch implantation, the switch is opened, and blood samples are collected 12 and 48 hours after switch opening. **e)** Quantification of Creatinine, BUN, sodium, and potassium before, 12 and 48 hours after VPS opening. The upper limit of the potassium assay is 9 mmol/L. **f)** Schematic diagram of uremia clearance. 24 hours after implantation, the switch was opened for 20 and then closed for 48 hours. Blood samples were collected before, 6, 12, 24, and 48 hours after VPS closure. Six VPS animals were operated and divided into two groups of three for blood collection (n=3 for 6 and 24h, n=3 for 12 and 48h) to reduce the number of repeated blood sampling for each mouse. **g)** Quantification of creatinine, BUN, sodium, and potassium before and 6, 12, 24 and 48 hours after VPS closure. **h)** Quantification of UPLC-M/MS-detected uremic toxins and amino acids in serum after 48h open VPS. *Statistics: For a statistical difference in mortality rates between CKD stages were calculated by univariate Cox regression. For e and g a two-way ANOVA (Tukey corrected) was calculated. For h a Welch‘s ttest was performed*.

Inspired by case reports of pseudo-kidney failure in patients with vesico-pertioneal fistulas, we aimed to induce uremia by generating a vesico-peritoneal shunt (VPS) to allow for peritoneal absorption of urinary solutes ^10^. We developed a method for the surgical implantation of a catheter into the bladder, which drains urine into the abdominal cavity (**Fig. S1d**). To enable discrete control of urine flow, we additionally engineered a bistable switch using a magnetic ball valve, which can be operated using an external magnet (**Fig. 1c, Fig. S1c**). Connection of the switch to the implanted VPS enables opening and closure of the VPS. After implantation we measured serum renal retention parameters and electrolytes after 12 and 48 hours of an open VPS (**Fig. 1d**). As a control, we performed VPS surgery but left the VPS closed during the experiment (sham). As expected, creatinine and BUN serum concentrations progressively increased in VPS mice over 12 and 48 hours culminating in an average BUN of 262 mg/dL at 48 h (**Fig. 1e**). Analogous to ESKD patients, VPS mice developed severe hyperkalemia (**Fig. 1e**)^11^. Further, VPS mice showed progressive hyponatremia, suggesting dilution due to fluid overload (**Fig 1e**). Abdominal sonography confirmed accumulation of intraabdominal urine with an open VPS, but not in sham mice (**Fig. S1 e-f**). Quantification of the intra-abdominal fluid revealed an average accumulation of 2.5 g urine over 48 hours with an open VPS (**Fig. S1g**). To determine whether our model permits for normalization of renal retention and electrolyte imbalance, we closed the VPS after an open interval of 20 hours and measured renal retention parameters and electrolytes at 6, 12, 24, and 48 hours (**Fig. 1f**). Immediately prior to closing the VPS, all animals displayed increased BUN and creatinine serum levels as well as hyperkalemia and hyponatremia (**Fig. 1g**). Closure of the VPS led to rapid normalization of creatinine and potassium levels. Similarly, serum concentrations of BUN and sodium returned to baseline and no significant differences between VPS and sham could be detected at 24 to 48 hours after switch closure (**Fig. 1g**). In summary, implantation of the VPS enables dialysis-like induction and clearance of uremia and electrolyte imbalance.

Kidney failure and dialysis is marked by a complex metabolic phenotype with accumulation of a variety of toxic water-soluble and protein-bound metabolites (uremic toxins), as well as dysregulation of amino acid handling ^12,13^. To test whether our model recapitulates uremia and metabolic dysfunction as seen in ESKD patients, we quantified a panel of uremic toxins and amino acids in serum by performing Ultra-Performance Liquid Chromatography coupled with Mass Spectrometry (UPLC-M/MS) (**Supp. Table 1**). After 48 hours of an open VPS, mice showed increased levels of Trimethylamin-N-oxid (TMAO), p-Cresylsulfate and Indoxyl-sulfate, representing disease-defining water-soluble (TMAO) and protein-bound (p-Cresylsulfate, Indoxyl-sulfate) uremic toxins in dialysis patients (**Fig. 1h**) ^14–16^. Analysis of amino acid metabolism revealed increased Kynurenine/Tryptophane-ratios and decreased Tyrosine/Phenylalanine-ratio in VPS mice, pointing towards an ESKD-characteristic metabolism of enhanced tryptophan degradation with kynurenine retention, and impaired conversion of phenylalanine to tyrosine (**Fig. 1h**) ^17,18^. Closure of the VPS rescued this phenotype, mirroring toxin-clearance and metabolic recovery as seen during dialysis (**Fig. S1h**). In summary, the VPS-model recapitulates metabolic dysfunction with intermittent accumulation of ESKD-defining uremic toxins and impairment of aromatic amino acid handling.

To assess performance of our model over time in a sex-balanced cohort we performed VPS implantation in male and female mice and subjected the latter to four weeks of cyclic uremia with 20 hours of an open and subsequent 28 hours of a closed VPS in repetitive cycles (**Fig. 2a**). Analogous to weight changes due to fluid overload in dialysis patients, VPS animals exhibited pronounced weight fluctuations, with an average weight gain of 1,5 g after 20 hours VPS opening, and an average weight loss of 1,1 g after 28 hours VPS closure **(Fig. 2b-c**). Overall, VPS animals showed an average increase in body weight of 11,5 % after 28 days (**Fig. 2d**). Next, we questioned whether intraabdominal fluid accumulation leads to systemic extravascular hypervolemia. In this regard, we quantified lung water and lung wet/dry ratios, a bona-fide marker of congestion and hypervolemia at 28 days after cyclic uremia. Indeed, VPS animals showed increased total lung water and an increased lung wet/dry ratio, suggesting the development of pulmonary edema in the context of extracellular hypervolemia (**Fig. 2e-f**). Confirming ESKD-defining uremia and electrolyte shifts, VPS mice showed increased serum BUN and creatinine levels as well as pronounced hyperkalemia and hyponatremia 2 and 4 weeks after surgery (**Fig. 2g-j**). Importantly, fluid retention, uremia and electrolyte dysbalance were observed independently of sex, underscoring applicability of the VPS model for sex-balanced studies of cyclic uremia.

**Figure 2:**
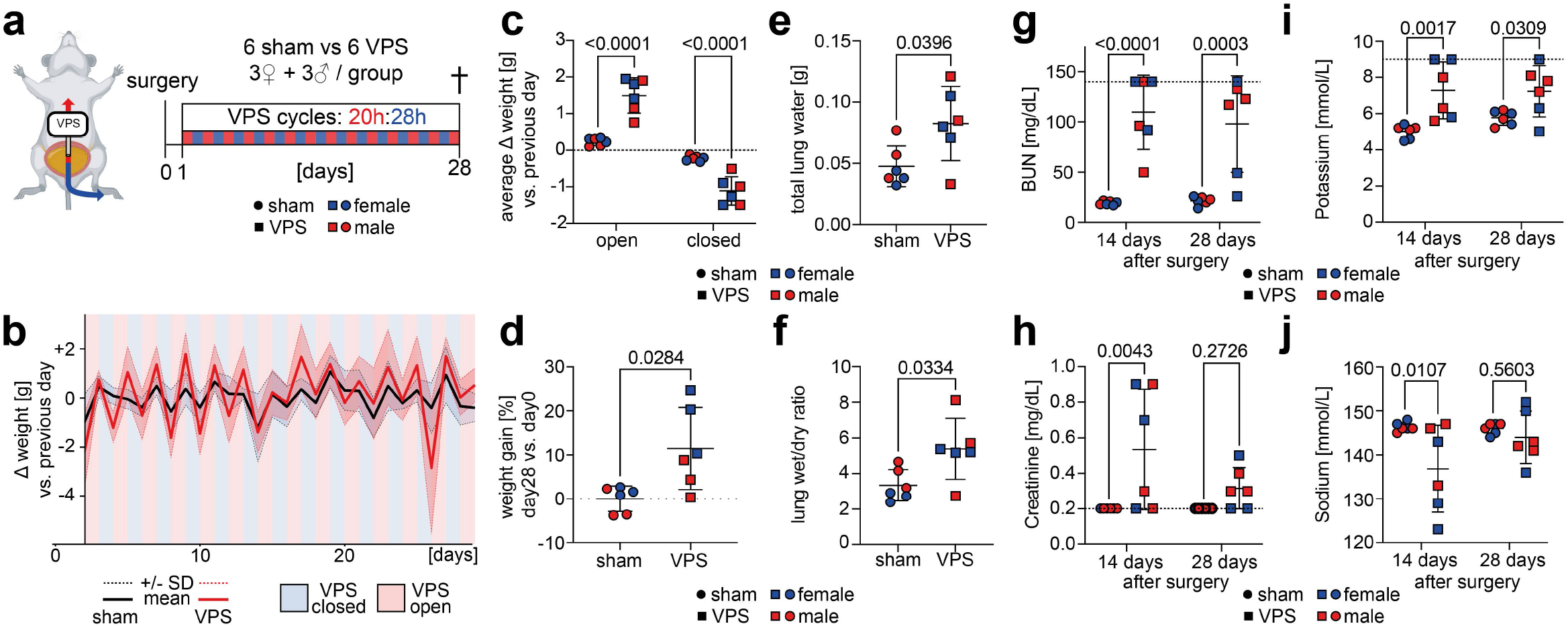
VPS mimics dialysis in a sex-balanced cohort. **a)** Schematic of the VPS time course. Uremia and dialysis were alternated as follows: 20 hours open VPS, 28 hours closed VPS. Mice were euthanized after 1 month. **b)** Line chart showing the average weight difference per mouse compared to the previous day over time, stratified by intervention group over time. **c)** Average change in animal weight 20 hours after opening and 28 hours after closing the magnetic switch, stratified by intervention group. **d)** Relative weight change 28 days after the start of the experiment, compared to individual starting weights. **e,f)** Total lung water and wet-to-dry ratio stratified by intervention group. **g,h)** Quantification of creatinine and BUN 14 and 28 days after surgery. The upper limit of the BUN assay is 140 mg/dL, while the lower limit of the creatinine assay is 0.2 mg/dL. **i,j)** Quantification of serum potassium and sodium levels 14 and 28 days after surgery. The upper limit of the potassium assay is 9 mmol/L. *Statistics: For c-f a Welch test was performed. For g-j a two-way ANOVA (Tukey corrected) was calculated*.

Next, we questioned whether VPS-induced cyclic uremia represents an independent driver of CVD in the presence of otherwise healthy kidneys. We performed VPS-implantation and investigated cardiac and immune cell dysfunction after 6 weeks of cyclic uremia (**Fig. 3a**). Measurements of BUN, creatinine, potassium, and sodium at 2, 4, and 6 weeks confirmed successful cyclic uremia over 6 weeks (**Fig. S2a**). To assess blood pressure in VPS mice, we performed volume pressure recording (VPR) during the uremic interval. Interestingly, VPS mice displayed decreased blood pressures compared to sham mice (**Fig. S2b**). To evaluate activity of the renin-angiotensine-aldosterone system (RAAS), we measured serum angiotensin II (AT2). In line with increased RAAS activity in ESKD, VPS mice showed increased AT2 levels in comparison to sham mice (**Fig. S2c**). In ESKD patients performing peritoneal dialysis, intraperitoneal instillation of hyperosmolar dialysates can lead to hypotension due to volume shifts into the abdominal cavity ^19,20^. Hypothesizing that hypotension in VPS mice is a consequence of hyperosmolar intraperitoneal urine, we measured osmolality and solutes in the intraperitoneal urine. As expected, intraabdominal fluid osmolality was higher than serum osmolality, indicating that hypotension may be a consequence of osmolality driven volume shifts into the abdominal cavity (**Fig. S2d**).

**Figure 3:**
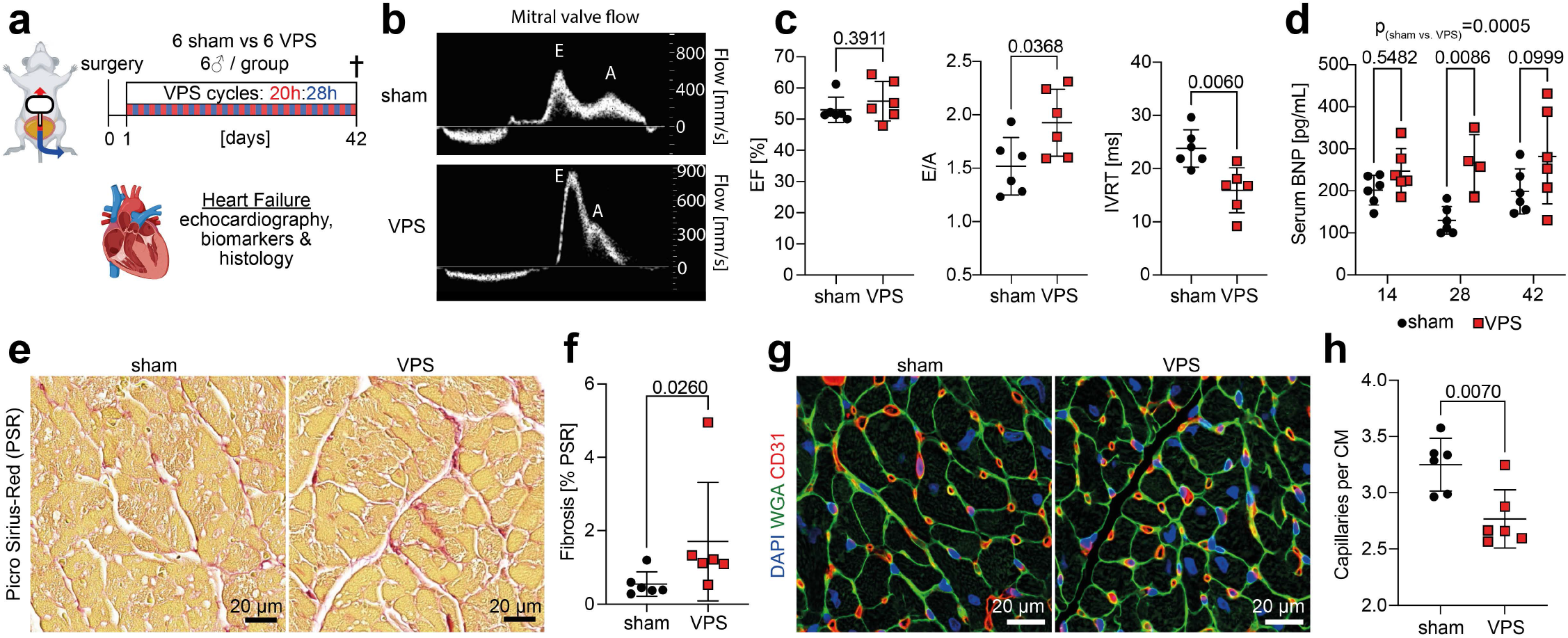
VPS-induced cyclic uremia drives heart failure. **a)** Schematic of the VPS timecourse over 6 weeks with cardiac readouts. Uremia and dialysis were alternated as following: 20 hours open VPS, 28 hours closed VPS. Mice were euthanized after 6 weeks. **b)** Representative images of echocardiographic pulsed-wave doppler of the mitral valve. **c)** Quantification of left ventricular ejection fraction (LVEF), E/A-ratio and isovolumetric relaxation time (IVRT) in sham and VPS animals after 6 weeks of cyclic uremia. **d)** ELISA quantification of murine brain natriuretic peptide (BNP) in serum after 2, 4 and 6 weeks of cyclic uremia. 2-way analysis revealed a significant increase of BNP in VPS mice over all timepoints (p=0.0005). **e)** Representative images of Picro Sirius-Red staining (PSR) in FFPE heart sections after 6 weeks cyclic uremia **f)** Quantification of left ventricular fibrosis as defined by PSR positive extracellular matrix in FFPE heart sections after 6 weeks of cyclic uremia. **g)** Representative images of Immunofluorescence for DAPI, Wheat Germ Agglutinin (WGA) and CD31 in FFPE heart sections after 6 weeks. **h)** Quantification of CD31^+^ capillaries per WGA+ cardiomyocyte (CM) in FFPE heart sections after 6 weeks of cyclic uremia. *Statistics: For c and h a Welch‘s Test was calculated. For d a two-way ANOVA (Tukey corrected) was calculated. For f a Mann-Whitney test was calculated*.

To investigate whether cyclic uremia induces heart failure, we performed echocardiography 6 weeks after VPS implantation (**Fig. 3b-c**). Quantification of the left ventricular ejection fraction (LV-EF) revealed no differences between sham and VPS animals (**Fig. 3c**). Sham and VPS mice showed comparable left-ventricular end-diastolic volumes (LV-EDV) and cardiac outputs (CO) (**Fig. S2e**). However, VPS animals exhibited an increased E/A ratio and a shortened isovolumetric relaxation time, key features of diastolic dysfunction in restrictive cardiomyopathy (**Fig. 3c**). Next, we quantified serum brain natriuretic peptide (BNP) levels, a key biomarker of heart failure, that also progressively accumulates with declining kidney function^21^. Indeed, serum BNP levels were increased in VPS compared to sham mice (main effect across timepoints with p=0.0005) (**Fig. 3d**). HFpEF and ESKD-induced heart failure are marked by interstitial fibrosis, capillary rarefaction and cardiomyocyte hypertrophy, representing the histological hallmarks of cardiac remodelling. Indeed, quantification of cardiac Picro Sirius-Red stainings revealed increased ECM accumulation in VPS animals in line with induction of cardiac fibrosis (**Fig. 3e-f**, **S2f**). Quantification of CD31^+^-capillaries revealed reduced capillary density in VPS animals, confirming capillary rarefaction (**Fig. 3g-h**). Notably, we did not observe cardiomyocyte hypertrophy. In line with the development of hypotension, the cardiomyocyte diameter was smaller in VPS mice compared to sham mice (**Fig. S2g**). Finally, as hyperphosphatemia is a hallmark of ESKD and an independent risk factor for CVD, we measured serum inorganic phosphate, which was significantly increased in VPS mice (**Fig. S2h**) ^22^. In summary, the VPS model allows for the exploration of uremia-induced heart failure characterized by echocardiographic signs of HFpEF, elevated levels of circulating HF biomarkers, and cardiac remodeling with fibrosis and capillary rarefaction, regardless of the presence of arterial hypertension.

A core feature of ESKD and CVD is immune dysfunction with loss of the adaptive immune response and parallel proinflammatory myeloid activation, leading to increased mortality rates in the context of infectious diseases ^23,24^. Blood count analysis 4 weeks after VPS implantation showed no differences in white blood cell count, red blood cell count, or hemoglobin between VPS and sham mice, with only moderately increased platelet counts in VPS mice (**Fig. S2i)**. Next, we assessed immune cell activation by performing single-cell RNA sequencing (scRNA-Seq) of CD45^+^ peripheral immune cells after 6 weeks of cyclic uremia. After preprocessing (demultiplexing, exclusion of doublets and low quality cells), we obtained 28,087 single cells from 11 mice (6 sham, 5 VPS) (**Fig. S3a-f**). Clustering based on marker genes identified eight major immune cell clusters (**Fig. 4a, S3g, Supp. Table 2**). Compositional analysis revealed VPS animals to exhibit reduced cell counts of *Ly6c2*^low^ monocytes and NK-cells, and, notably, nearly complete depletion of B-Cells (**Fig. 4b-c**, **Fig. S3h**). *MultiNicheNet* ligand-receptor analysis revealed increased signaling from granulocytes and monocytes via Amyloid β precursor protein (*App*) to CD74 on B-cells, recently described to drive B-cell suppression in sepsis ^25,26^ (**Fig. 4d**). Next, we aimed to investigate *Ly6c2*^high^ monocytes representing widely recognized orchestrators of uremia-driven immune dysfunction. *PROGENy* pathway analysis revealed increased activity of proinflammatory (NF-κB and TNFα) and profibrotic (TGFβ) pathways in *Ly6c2*^high^ monocytes (**Fig. 4e**). Importantly, analysis of differentially expressed genes in *Ly6c2*^high^ monocytes revealed increased expression of established proinflammatory and proatherogenic CKD-biomarkers such as calprotectin (*S100a8/9*), mincle (*Clec4e*) as well as profibrotic molecules including lipocalin-2 (*Lcn2*) and transforming growth factor-beta-induced (*Tgfbi*) (**Fig. 4f, Supp. Table 3**) ^27–32^. Geneset Enrichment Analysis (GSEA) for Gene Ontology: Biological Processes (GO:BP) revealed enrichment of terms associated with leukocyte migration and inflammatory response among upregulated genes and enrichment of terms associated with cytokine production among downregulated genes in VPS compared to sham animals (**Fig. 4g**). The simultaneous upregulation of proinflammatory genes alongside anti-inflammatory genes such as *Arg2* underscores the paradoxical phenomenon of immune activation with parallel immune paralysis in ESKD-immune dysfunction, that has been more recently linked to ESKD-induced immunosenescence ^24,33–35^. Accordingly, we used *SenePy,* a database of cell-specific senescence gene signatures, to evaluate uremia-induced senescence in immune cells ^36^. Indeed, senescence scoring confirmed VPS-induced upregulation of immunosenescence defining genes in monocytes with strongest increase in *Ly6c2*^high^ monocytes (**Fig. 4h, Fig. S3i**). Importantly, *Ly6c2*^high^ monocytes also showed increased expression of NABA ECM regulator genes, which we recently demonstrated to be a marker for maladaptive profibrotic macrophages (**Fig. 4i, Fig. S3j**) ^37,38^. In summary, cyclic uremia in the VPS model mimics ESKD-defining immunosenescence characterized by B-cell depletion, increased proinflammatory and profibrotic activation as well as enrichment of pro-migratory genesets in monocytes.

**Figure 4:**
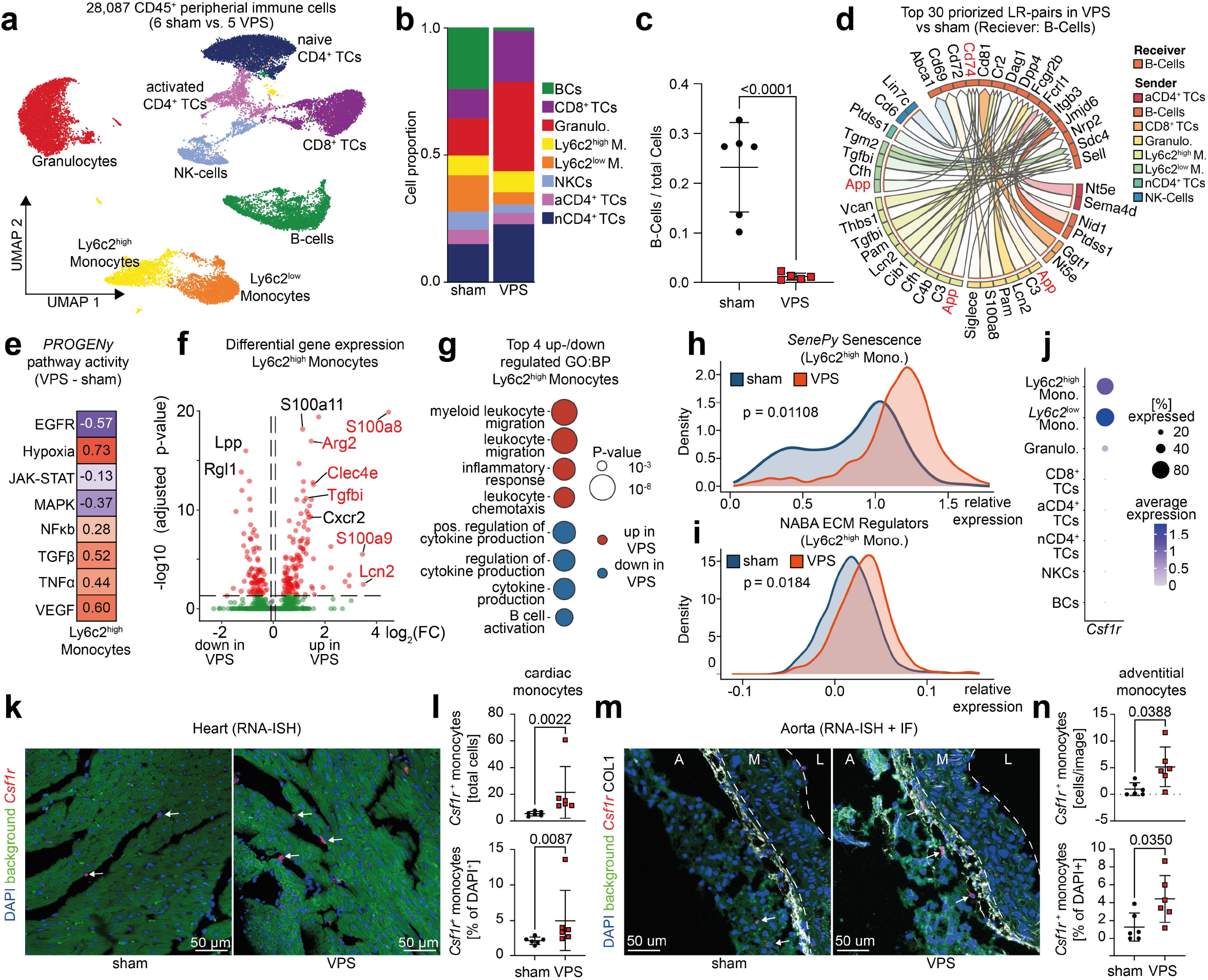
VPS induces ESKD-associated immunosenescence and drives mononuclear chemotaxis. **a)** UMAP embedding of 28,087 high quality single cells from scRNAseq of peripheral CD45^+^ immune cells in 5 VPS and 6 sham mice after 6 weeks of VPS-modeling (20 h open; 28 h closed). **b)** Barplot showing the cellular composition stratified by intervention. **c)** Scatter dot plot of B-Cells as a fraction of total peripheral immune cells per mouse stratified by intervention. **d)** Top 30 VPS-specific MultiNicheNet Ligand-Receptor Pairs with B-Cells as receivers sorted by prioritization score. **e)** Heatmap of differential PROGENy pathway activity (VPS vs. sham) in *Ly6c2*^high^ Monocytes. **f)** Volcano Plot of pseudobulk differential gene expression analysis (DESeq2) in *Ly6c2*^high^ Monocytes. Red dots indicate genes that are significantly up-/downregulated. **g)** Dotplot with top 4 enrichedGO:BP terms based on up-and downregulated genes (VPS vs. sham) in *Ly6c2*^high^ Monocytes. Terms with a term size >800 genes were excluded. **h)** Density plot showing relative expression of the SenePy senescence gene signature in *Ly6c2*^high^ Monocytes. **i)** Density plot showing relative expression of the NABA ECM regulator genes in *Ly6c2*^high^ Monocytes. **j)** Dotplot showing the expression of Csf1r across all immune cell clusters. **k)** Representative confocal images of RNA-ISH for Csf1r in heart sections. **l)** Quantification of absolute and relative (per DAPI^+^ cells) Csf1r^+^ monocytes in heart sections. Arrows point towards Csf1r^+^ monocytes. **m)** Representative confocal images of RNA-ISH for Csf1r and IF for COL1 in aortic sections. A: adventitia, M: media, L: lumen; Arrows point towards Csf1r^+^ monocytes. **n)** Quantification of Csf1r^+^ monocytes in the adventitial layer in aortae. *Statistics: For b per-animal cell-type proportions were compared between sham and VP-shunt using a robust empirical Bayes moderated test on arcsine-transformed proportions (propeller). For f DESeq2-integrated Wald-Test was performed on pseudobulked data and FDR-corrected p-values were used. For h and i a two-tailed t-test was performed. For k a Mann-Whitney test was performed*.

To validate our scRNAseq findings of an increased monocyte activation, we lastly assessed monocyte infiltration in heart and aortas. RNA-In-situ-hybridization (RNA-ISH) for *Csf1r*, a bona fide marker of mononuclear phagocytes, confirmed increased monocyte accumulation in VPS hearts in comparison to sham hearts (**Fig. 4j-l**). To assess whether aberrant monocyte expansion extends beyond the heart, we also performed RNA-ISH for *Csf1r^+^* monocytes in aortae of VPS and sham mice (**Fig. 4m**). Confirming our findings in the heart, *Csf1r^+^* monocytes were enriched in the adventitial tissue of aortas in VPS compared to sham (**Fig. 4n**).

In summary, we demonstrate that the VPS model drives an immunosenescent monocyte phenotype with proinflammatory, profibrotic *Ly6c2*^high^ monocyte activation and increased monocyte expansion in hearts and aortas.

Dialysis patients represent a high-risk CVD group, for which therapeutic interventions are lacking ^39^. Currently, translational research and drug development efforts are constrained by the shortage of clinically relevant mouse models for these patients ^39^. Importantly, existing rodent CKD-models do not resolve the effects of uremia from well-established drivers of CVD such as hypertension and ultimately fail to recapitulate CKD-driven CVD ^7–9^. Here, we present an easy-to-establish, sex-inclusive mouse model for reversible, on-demand induction and clearance of uremia. Critically, our model for the first time uncouples the kidney’s endocrine (synthesis of i.e., erythropoietin, vitamin D, renin) and excretory functions, two systems that are otherwise considered inseparable in established CKD models. Iterative application of our model enables induction of cyclic uremia recapitulating ESKD-like metabolic shifts with cyclic accumulation of uremic toxins, impaired amino acid handling and electrolyte imbalance. While we focus here on cyclic uremia as a proof-of-principle application, the VPS is not restricted to this setting and may be leveraged to dissect uremia in other conditions, such as AKI or combined endocrine and exocrine kidney failure (⅚ nephrectomy + VPS).

At the metabolic level, accumulation of toxic metabolites is a hallmark of ESKD-driven cardiovascular disease. TMAO, p-Cresylsulfate and Indoxyl sulfate are both drivers and biomarkers for cardiovascular complications such as myocardial infarction, atherosclerosis and heart failure ^16,40–44^. Existing mouse models studying uremic toxin-induced effects are largely based on dietary supplementation (e.g., phosphate or adenine) and therefore limited in their pathophysiological relevance ^45^. In contrast, uremia in VPS mice is caused by intermittent uncoupling of kidney clearance leading to accumulation of cardiotoxic metabolites without dietary modifications. This resolves a fundamental gap in the field of uremia research, as the VPS model captures the cyclic accumulation and fluctuation of a broad spectrum of uremic toxins, enabling study of their synergistic toxicity without confounding from a dysfunctional renal endocrine system or off-target toxicity from unphysiologic dietary supplementation.

HFpEF represents the predominant manifestation of CKD-driven heart disease ^46^. In line with the clinical criteria for diagnosis of HFpEF - (i) symptoms of heart failure, (ii) structural or functional signs of cardiac dysfunction with (iii) an LV-EF > 50% ^47^ - we demonstrate that the VPS model leads to pulmonary edema, elevated BNP levels and echocardiographic signs of diastolic dysfunction. At the histological level, we confirm structural cardiac remodelling in VPS mice defined by increased fibrosis and capillary rarefaction. Intriguingly, while in-vivo effects of uremia have not yet been studied separately from CKD-induced arterial hypertension, we are able to show development of HFpEF to be independently driven by a uremic metabolism.

Infections are the second leading cause of death among dialysis patients and can be attributed to reduced pathogen defense due to ESKD-induced immunosenescence ^48^. In line with the recent association of B-cell lymphopenia in ESKD patients, VPS modeling led to B-cell depletion in mice ^49–51^. In addition, we find that cyclic uremia induces a proinflammatory and profibrotic activation of monocytes, marked by increased expression of calprotectin (*S100a8/a9*) and Mincle (*Clec4e*), both well-defined markers for CVD and fibrosis ^27,28,30,52^. Finally, we link proinflammatory monocyte activation to cardiac remodelling and vascular inflammation by demonstrating that cyclic uremia and immunosenescence results in increased monocyte infiltration into the heart and vasculature. Further analysis of local immune niches in VPS mice offers a wide range of opportunities to investigate disease-specific mechanisms of the maladaptive immune response in ESKD.

In summary, we present the first preclinical mouse model of uremia that enables investigation of the effects of a broad spectrum of uremic toxins independent of endocrine dysfunction of the kidneys. Long-term application of our model with iterative cycles of uremia recapitulates ESKD-defining metabolism. Strikingly, we identify cyclic uremia as an independent driver of CVD-defining hallmarks such as immunosenescence, vascular inflammation and HFpEF in the presence of otherwise healthy kidneys. We demonstrate that cyclic uremia leads to proinflammatory monocyte activation and migration in the absence of tissue insults, linking immunosenescence and tissue remodeling in ESKD. As a resource, we envision that the easy-to-establish, sex-balanced VPS model will fill a critical translational gap for interrogation of ESKD-driven CVD *in vivo*.

## Methods

### Mice

All animal experiments were approved by the regional authorities (LANUV-NRW, Germany; Animal Welfare). 8 - 14 weeks old 129/Sv mice were housed in cages with up to four other animals under pathogen-free conditions with a 12-hour day-night cycle. Water and standard food were provided ad libitum. The room temperature was maintained at a constant 20°C.

### Construction of a magnetic VPS switch

The components were designed using CAD Inventor Professional and milled on a DMG Mori 5-axis CNC milling machine. The sealing rings used were made from silicone supplied by Wacker. The screws, dowel pins, and ball are made of corrosion-resistant steel. The switch consists of an upper and a lower component (10×8×2 mm) made of Susta Peek MG blue BL. The upper component features a guide channel for the ferromagnetic bead, a groove for a sealing ring, two alignment holes for positioning the lower component, and four countersunk holes for M1 cylinder head screws. The lower component consists of a borehole for the inflow and outflow of urine, a circular pocket for a sealing ring, two alignment bores, four M1 tapped holes for positioning and securing the upper component, and two circular pockets for the magnets. The switch was installed as follows: First, two locating pins are installed in the designated positions on the lower component, and the sealing ring for the left bead position (valve closed) is inserted. On the opposite side of the component, two magnets are glued into the designated circular pockets (with both magnets oriented in the same direction). In the upper component, the sealing ring is inserted into the designated groove and the ferromagnetic bead is inserted into the designated guide channel. Both components are connected using four M1 screws; the dowel pins installed in the lower component and the pilot holes in the upper component ensure precise positioning. The screws are tightened using a torque wrench. The switch has final dimensions of 10×5×8 mm³.

### Surgical implantation of the VPS

Mice were anesthetized via intraperitoneal administration of a combination anesthetic (ketamine and xylazine, dissolved in 0.9% saline) at a dose of 120 mg/kg body weight ketamine + 16 mg/kg body weight xylazine and placed on a heated warming pad (37° Celsius) to maintain body temperature. Metamizole (200 mg/kg body weight) was administered subcutaneously for analgesia. The abdomen was shaved and disinfected with povidone-iodine. The peritoneal cavity was opened via laparotomy, and the urinary bladder was exposed and fixated with medical tape containing an opening for the bladder. After placing a suture (Ethilon 7-0), a puncture was made at the bladder dome and an 18G catheter (FEP-Teflon) was inserted using the Seldinger technique. The catheter was secured using the preplaced suture. The (closed) VPS switch was connected to the cranial, extravesical end of the catheter, secured to the catheter with tissue adhesive (Histoacryl), and placed intraperitoneally. The peritoneum and the skin were closed with an interrupted suture (Mersilene 3-0). Until awakening, heat was applied using an infrared lamp under supervision. For further analgesia, metamizole (200 mg/kg body weight based on an estimated water intake of 4 ml per animal per day) was administered via the drinking water until the third postoperative day. From the first preoperative day until the end of the experiment, enrofloxacin (0.25 mg/mL) was administered via the drinking water to prevent infection. In long-term experiments, the mouse weight was measured daily starting from the first time the switch was opened.

### Induction and clearance of uremia

Using an external magnet, the ferromagnetic bead can be moved transdermally within the guide channel of the upper component between two positions (left and right). After the external magnet is removed, the ball remains on the desired left (VPS valve closed) or right (VPS valve open) side of the guide channel, where it is being held in place by one of two magnets located approximately 2 mm below it in the lower component. Bead position on the right side: the valve is open, enabling urine flow flow from the bladder through the inlet port (supply line) via the ball’s guide channel to the outlet port (abdominal cavity). Bead position on the left side of the guide channel: The ball is drawn into the silicone sealing ring by the magnet located in the downer compartment and in consequence blocks urine flow from the bladder into the abdominal cavity. When operating the VPS in mice, mice were held by the scruff of the neck, and the magnet was placed against its abdomen. The switch was held in place transdermally with the fingertips and was then opened or closed by moving the magnet to the left or right side, respectively. In sham animals, the switch was left in the closed position. For long term experiments the VPS was opened for 20 h and closed for 28 h.

### Mortality analysis in the UK biobank

We conducted a retrospective cohort analysis using linked UK Biobank hospital inpatient records and mortality registry data. CKD was defined based on staged ICD-10 codes N18.1–N18.5. For each participant with staged CKD, the index date was defined as the first recorded staged CKD diagnosis, and CKD stage was assigned according to the corresponding ICD-10 code at this first staged diagnosis. Individuals with kidney transplant codes Z94.0 or T86.1 were excluded. A mutually exclusive non-CKD comparison group was constructed. Participants with any staged CKD code N18.1-N18.5 at any time during available hospital follow-up were assigned exclusively to the CKD cohort and were not eligible for the control group. Controls were therefore defined as individuals without any staged CKD diagnosis during follow-up. Other kidney-related or unspecified renal ICD-10 codes were not used to exclude controls but were retained as descriptive quality-control variables. Follow-up for controls started at the mean CKD index date observed among CKD participants. Follow-up started at the first staged CKD diagnosis for CKD participants. Participants were followed until death or administrative censoring on 25 August 2023, whichever occurred first. Individuals with death before the respective baseline date were excluded from the risk set. All-cause mortality was defined using registry-derived date of death. Cause-specific mortality was classified according to the primary ICD-10 cause of death and grouped into cardiovascular, cancer, respiratory, infectious, digestive/liver, renal, and other or unknown mortality. For descriptive analyses, we calculated participant numbers, deaths, person-years, and mortality rates per 100 person-years for controls and CKD stages G1-G5. To assess the association between CKD stage and mortality, univariate Cox proportional hazards models were fitted for all-cause and cause-specific mortality endpoints.

### Blood collection and analysis

Animals were anesthetized with isoflurane (2–2.5 % (v/v)). 80 µl of blood were collected from the retrobulbar venous plexus. Hemoglobin, serum electrolytes (sodium, potassium), and renal retention values (creatinine, BUN) were measured with a CHEM8+ cartridge (Abbott) using the i-STAT 1 system (Abbott). The CHEM8+ cartridge has a lower limit for Creatinine (0.2 mg/dL) and upper limits for potassium (9 mmol/L) and BUN (140 mg/dL). The complete blood count (VPS 6 weeks) was determined by flow cytometry. For serum analysis, blood was stored in serum tubes (Sarstedt) for 30 minutes and the serum was separated from cellular components by centrifugation (2000 xG, 10 minutes).

### Quantification of BUN, BNP and phosphate

Serum was prepared as described above. BUN assay (Invitrogen, Catalog Number EIABUN), BNP assay (Invitrogen, Catalog Number EEL089) and phosphate assay (ab65622, abcam) were performed as colorimetric assays according to the manufacturer’s protocols. Due to low maximum blood sample volume for blood withdrawal in ongoing mouse experiments, BNP could not be quantified in 2 mice at 4 weeks (VPS) and phosphate could not be quantified for one mouse (VPS).

### Lung water quantification

The lungs of the animals were removed at the hilum immediately after euthanasia, prior to perfusion. The weight (wet weight) was measured immediately after euthanasia. The lungs were then dried at 80 °C for 48 hours and the dry weight was subsequently determined.

### Serum mass spectrometry

Serum was processed as described above and a panel of uremic retention solutes was analyzed using ultra performance liquid chromatography coupled with tandem mass spectrometry (UPLC-M/MS) (Acquity – Xevo TQS, Waters, Zellik, Belgium), as described previously ^53^. Briefly, 50 µl serum sample, 20 µl of internal standard mixture and 200 µl acetonitrile were thoroughly mixed in 96-well Ostro plates (Waters, Zellik, Belgium). After separation by positive pressure manifold, the organic phase was removed by a gentle stream of nitrogen for 30 minutes at 40°C and dissolved with 1000 µl of Milli-Q water; 5 µl of the final solution was injected on the UPLC-MS/MS system. Chromatographic separation was performed on a Acquity CSHFluoroPhenyl column (50 x 2.5 mm; 1.7 µm particle size; Waters). The mobile phase, delivered at a flow rate of 0.5 ml/min at 40 °C, was a gradient of 0.1 % formic acid in Milli-Q water and methanol. Ionization was achieved using alternating electrospray positive ionization mode (ESI+) and negative ionization mode (ESI-). The multiple-reaction monitoring (MRM) transitions, cone voltage and collision energy were optimized for each individual compound. The total, within-run, between-run and between-day method imprecision according to the NCCLS EP5-T guideline were below 15% for all compounds. Mean recoveries were between 83 % and 104 % for all the compounds. Due to low blood sample volume, UPLC-MS/MS-quantification of metabolites could not be performed in one mouse (VPS).

### Single-cell RNA sequencing of immune cells

Immune cells were isolated from citrate-anticoagulated whole blood. First, cells were isolated by centrifugation (400 xG, 5 minutes, room temperature). Subsequently, cells were resuspended in erythrocyte lysis buffer (BD Bioscience, Catalog No. 555899) and incubated for 6 minutes at room temperature. The cell suspension was then diluted with MACS buffer (PBS, 0.5% BSA, 2 mM EDTA), filtered through a 30-µm strainer, and washed at 400 xg for 5 minutes at 4 °C. The cell pellet was resuspended in MACS buffer and incubated with Hashtag antibodies (Biolegend, TotalSeq™-A030X) and anti-CD45 microbeads (Miltenyi Biotec, 130-052-301) for 15 minutes at 4 °C. Following an additional wash step, magnetic separation of the CD45⁺ cells was performed using MS columns (Miltenyi Biotec) according to the manufacturer’s instructions. From the isolated cells, cDNA libraries were generated for single-cell RNA sequencing using the 10x Genomics platform (Chromium GEM-X Single Cell 3’ v4) according to the manufacturer’s protocol. The quality of the libraries was verified using a 2200 TapeStation system (Agilent Technologies). Sequencing of the cDNA libraries was performed on an Illumina NovaSeq system targeting 20,000 reads per cell.

### Analysis of single-cell RNA sequencing data

The data were aligned to the murine genome (GRCm39-2024-A) using *CellRanger* (version 9.0.1), and a count matrix was generated. Demultiplexing of individual animals was performed using the *Cite-Seq* protocol based on hashtag sequence expression ^54^. Due to the low expression of the hashtag sequence with resulting inability to demultiplex, one mouse (VPS) was removed from the dataset, resulting in 58,189 cells prior to QC. Further analyses were conducted using Seurat (v5.4) ^55^. Cells with >10% mitochondrial RNA, >40,000 detected transcripts and <200 or>60,000 Features counts were excluded. Doublets were detected using *scDblFindeR* and excluded based on the default class annotation (*class == ‘doublet’*) ^56^. The datasets were integrated using *harmony* (*k.param == 20*) and normalized (*LogNormalize, scale.factor = 10,000*) followed by z-score scaling (ScaleData) of all genes. Variable features were identified using the variance-stabilizing transformation (4,000 HVGs). Nearest-neighbor graphs were built on the *harmony* embedding and clusters were identified with the *Leiden* algorithm (*25 harmony dimensions, resolution 0.1*). Cluster annotation was performed manually based on the corresponding marker genes. Manual annotation detected 2 doublet-like clusters (clusters 8+9) with high counts of erythrocyte-specific genes that were excluded from downstream analysis, resulting in 8 final immune cell clusters with a total of 28,087 high-quality cells from 11 animals. For compositional analysis, the proportion of each cell type was calculated in each mouse (*HTO-classification*) individually, differences between VPS and sham animals were calculated using the *propeller* package (*transform = “asin”*) and significance was evaluated using propeller’s FDR-adjusted p-values ^57^. Differential gene expression was assessed separately for each annotated immune cell cluster using pseudobulk RNA-seq profiles generated by summing normalized counts per mouse and pairwise comparison with *DESeq2* ^58^. P-values were adjusted for FDR using the Benjamini–Hochberg correction. For each cluster, significant *DESeq2* genes were tested for over-representation against *GO Biological Process* using *g:Profiler* (gprofiler2::gost, organism *mus musculus*, significance threshold 0.05), with enriched terms filtered to gene set size ≤ 800 for visualization ^59^. Pathway activity scores were inferred at single-cell resolution using *PROGENy* (*organism == ‘mous’*) applied to the log-normalized expression matrix, then summarized as mean pathway activity per annotated cluster and condition ^60^. For differential pathway activity the average pathway activity scores of VPS and sham were subtracted. For ligand-receptor analysis we performed *MultiNicheNet* analysis and identified the top 30 prioritised ligand receptor pairs in sham and VPS mice with B-cells as receiver ^61^. For senescence-scoring we computed an expression score for each cell by aggregating weighted normalized expression values of *SenePy*-defined murine signature genes. Similarly, we computed NABA ECM regulator expression by computing a module score (*AddModuleScore)* for expression of the defined murine orthologs of NABA ECM regulator genes.

### Abdominal ultrasonography and echocardiography

The ultrasound examination was performed under isoflurane anesthesia (initially 5% (v/v)] in the anesthesia chamber, followed by 2–3% (v/v) via nasal mask). Anesthesia depth was controlled by monitoring heart rate and respiratory rate. To maintain body temperature, the animals were placed on a heating pad (37 °C). The thorax (echocardiography) or the abdomen (abdominal sonography) was depilated using depilatory cream. This was followed by abdominal sonography (to assess the position of the VPS and to detect ascites) and echocardiography using a small-animal ultrasound device (Vevo 3100 and MX550D transducer, FUJIFILM Visualsonics). Left ventricular ejection fraction was measured using Simpson’s method, while diastolic dysfunction was quantified by measuring peak doppler blood inflow velocity across the mitral valve during early (E) and late diastole (A), as well as isovolumetric relaxation time (IVRT). Analysis was performed by a trained professional using the VevoLab Software (FUJIFILM, VEVO LAB).

### Volume pressure recording (VPR) for blood pressure measurement

Non-invasive blood pressure measurements were performed by immobilizing non-anesthetized mice in polycarbonate tubes and determining the tail blood flow rate using a volume-pressure sensor and a tail cuff.

### Histological analysis of hearts

After euthanasia, the hearts were first perfused with PBS, and the cardiac base was fixed in 4% paraformaldehyde (PFA) in PBS for 24 hours and embedded in paraffin. 1 µm sections were prepared using a microtome. The sections were deparaffinized in xylene (3 × 5 minutes) and rehydrated using a descending ethanol series (100%–96%–96%–70%). Picro-Sirius Red (PSR) staining was performed using a PSR staining kit (Morphisto) according to the manufacturer’s protocol. The slide was imaged using a 40x objective in an Aperio AT2 Slide Scanner (Leica Biosystems). The images were analyzed in a blinded manner by a trained pathologist using Aperio ImageScope software. PSR-positive pixels were detected based on an intensity threshold for the red spectrum. The same threshold was used for all tissue samples. Total fibrosis was defined as the ratio of PSR-positive pixels to total pixels in the tissue.

### Immunofluorescence staining of heart sections

FFPE slides were deparaffinized, rehydrated, and heat-induced antigen retrieval was performed in citrate buffer. The slides were stained with fluorescein-conjugated WGA (wheat germ agglutinin; 1:50; FL-1021; Vector Laboratories; Burlingame, CA, USA; RRID:AB_2336866) and CD31 (Cluster of Differentiation 31; 1:50; AF3628; R&D Systems; Minneapolis; USA; RRID:AB_2161028). An anti-goat antibody (1:200; BA-5000; Vector Laboratories, Burlingame, CA, USA; RRID:AB_2336126) was used to detect CD31, and the VECTASTAIN Elite ABC-HRP Kit (Vector Laboratories; Burlingame, CA, USA) was used for amplification. Staining was developed with Opal 570 fluorophores (PerkinElmer Life and Analytical Sciences; Boston; MA; USA), and cell nuclei were counterstained with DAPI. Images of the left ventricular wall showing cross-sectioned cardiomyocytes were acquired at a resolution of 332 × 220 µm using a Zeiss Axio Imager 2 with a 40× objective and ApoTomePlus (Carl Zeiss AG, Oberkochen, Germany). The images were analyzed using an ImageJ pipeline, as described before ^62^.

### RNA in situ hybridization (RNA-ISH) in heart and aorta tissue

RNA-ISH was performed with the RNA-Scope detection kit (Multiplex Fluorescent Reagent Kit v2 Assay). The staining procedure was performed according to the manufacturer’s protocol for formalin fixed, paraffin embedded (FFPE) sections (heart) and formalin-fixed OCT-embedded sections (aorta). We performed target retrieval for 30 min at 99°C and reduced the protease-treatment to 10 min. For RNA-hybridization of *Csf1r* we used the probe C2-m*Csf1r* (Cat No. 428191-C2) and utilized Cy2 (heart) and Cy5 (aorta) as fluorescent dyes. For additional immunofluorescence staining for COL1 in aortae, tissues were blocked after RNA-ISH with 10% BSA for 30 min and incubated for 1 h at room temperature with primary antibody (Goat Anti-Type I Collagen, Cat. No.: 1310-01, Southernbiotech, 1:100) and stained with a secondary antibody (AF488-anti-Goat) for 30 min at room temperature. Counterstaining was performed with DAPI (1 μg/mL) for 2 min. Sections were mounted with ImmuMount mounting media, covered with coverslips and imaged on a confocal microscope (Nikon A1R confocal laser microscope, channels 405 nm, 488 nm, 564 nm and 647 nm (Cy5)). Analysis was performed using a standardized workflow with Max-IP Z-projection and automated adaption of brightness in ImageJ and consecutive cell detection and object classification in QuPath. Nuclei were segmented with the cell detection tool based on DAPI channel (threshold = 250(heart)/110(aorta), cell expansion = 3 μm (heart)/2 μm (aorta)). Object classification for *Csf1r* was then performed using the single measurement classifier for *Csf1r* signal. All images were processed as a batch and parameters were set equally ^63,64^.

## Statistical analysis

Data is shown as mean ± SD. To test statistical differences between two groups with gaussian distribution, a student t-test (homoscedastic distribution) or Welch two-tailed t-test (heteroscedastic distribution) was performed. To test statistical differences between two groups with non-gaussian distribution, a Mann-Whitney-Test was performed. To evaluate statistical differences between 2-factor sample data, a two-way ANOVA with Tukey correction for multiple testing was performed. Graph-Pad Prism (version 10) was used for statistical analysis. Statistical testing in scRNA-seq Data and UK-Biobank data is discussed in the corresponding methods sections. Given a p-value < 0.05, a difference was considered to be significant.

## Data and code availability

The original code is available on github under https://github.com/GiscSC/scRNA_ImmuneCells_6weeks.

ScRNA-Seq data has been deposited at Zenodo and is available under 10.5281/zenodo.21979287 using the following link: https://zenodo.org/records/21979287?preview=1&token=eyJhbGciOiJIUzUxMiJ9.eyJpZCI6IjdjZGVjN2I1LTVjNmItNDJkYy1iY2YxLTAxOWY5Mjg1MGE1MSIsImRhdGEiOnt9LCJyYW5kb20iOiJmOTMxZjYzOWU2ZDgyZWQyZjVjYzAwN2ExMzM3Mjc3YyJ9.dyPZLY7gUeMKeR-0F9cb3MGOD0b6waa2fmWTvWlHJKPsXZyUyPm-mBSGbcDHqk3yzr8SalRVjPZri1fgwymHeg

## Acknowledgments

This work was supported by the Excellence Strategy of the Federal Government and with support from the Exploratory Research Space (ERS) of RWTH to GJLS (EXS-SU-StUpPD_506-25). GJLS was further supported by a RWTH Aachen University Clinician Scientist grant as well as an RWTH Aachen University START grant (113/24). R.K. was supported by the German Research Foundation (DFG: SFBTRR219 322900939, CRU344 4288578857858, CRU5011 445703531, KR4073/18-1 (561848128)), the European Research Council (ERC-CoG 101043403, ERC-PoC 101212974), the BMBF consortia CureFib (BMBF 01EJ2201A and 01EJ2501A), and the BMBF consortia KMUi DecodeFibrosis (BMBF 16LW0578). This work was supported by the Transgenic Core Facility, a core facility of the Interdisciplinary Centre for Clinical Research (IZKF) Aachen within the Faculty of Medicine at RWTH Aachen University. PB is supported by the DFG (Project IDs 322900939 & 445703531 & 552234081) and the European Research Council (ERC-CoG 101001791). This research has been conducted using the UK Biobank Resource under Application Number 71300. UK biobank data was accessed by C.V.S and K.M.S. Copyright © 2026, NHS England. Re-used with the permission of the NHS England and/or UK Biobank. All rights reserved. This work uses data provided by patients and collected by the NHS as part of their care and support.

## Authors contributions

KH, GJLS, AH and RK initiated the study. GJLS wrote the manuscript and organized the figures. AH, KH and GJLS designed the VPS switch. GJLS established and performed the surgery with assistance from KH, CS, XL, SM, LK and NL. GJLS and KH performed animal experiments with assistance from CS, XL, MK, NL, LK, LL, QL, KV, ASA, AB and DS. GJLS performed scRNASeq analysis. KH and GJLS analyzed the echocardiography data. PD, GJLS and PB performed histological stainings and analysis. JdL and BM performed UPLC. XL and GJLS performed RNA-ISH in heart and aorta. CVS and KMS performed mortality analysis of the UK biobank register. All authors have read and accepted the manuscript.

## Declaration of generative AI and AI-assisted technologies in the writing process

*Cursor* (Anysphere Inc.) was used to improve the readability and consistency of the R-based scripts and repositories. *OpenEvidence* was used to diversify literature research. *DeepL* and *ChatGPT* were used to assist with language editing and improve the readability of the manuscript. The authors reviewed and edited the output as needed and take full responsibility for the content of the published article.

## Supplemental Figures and Supplemental Figure Legends

**Figure S1:**
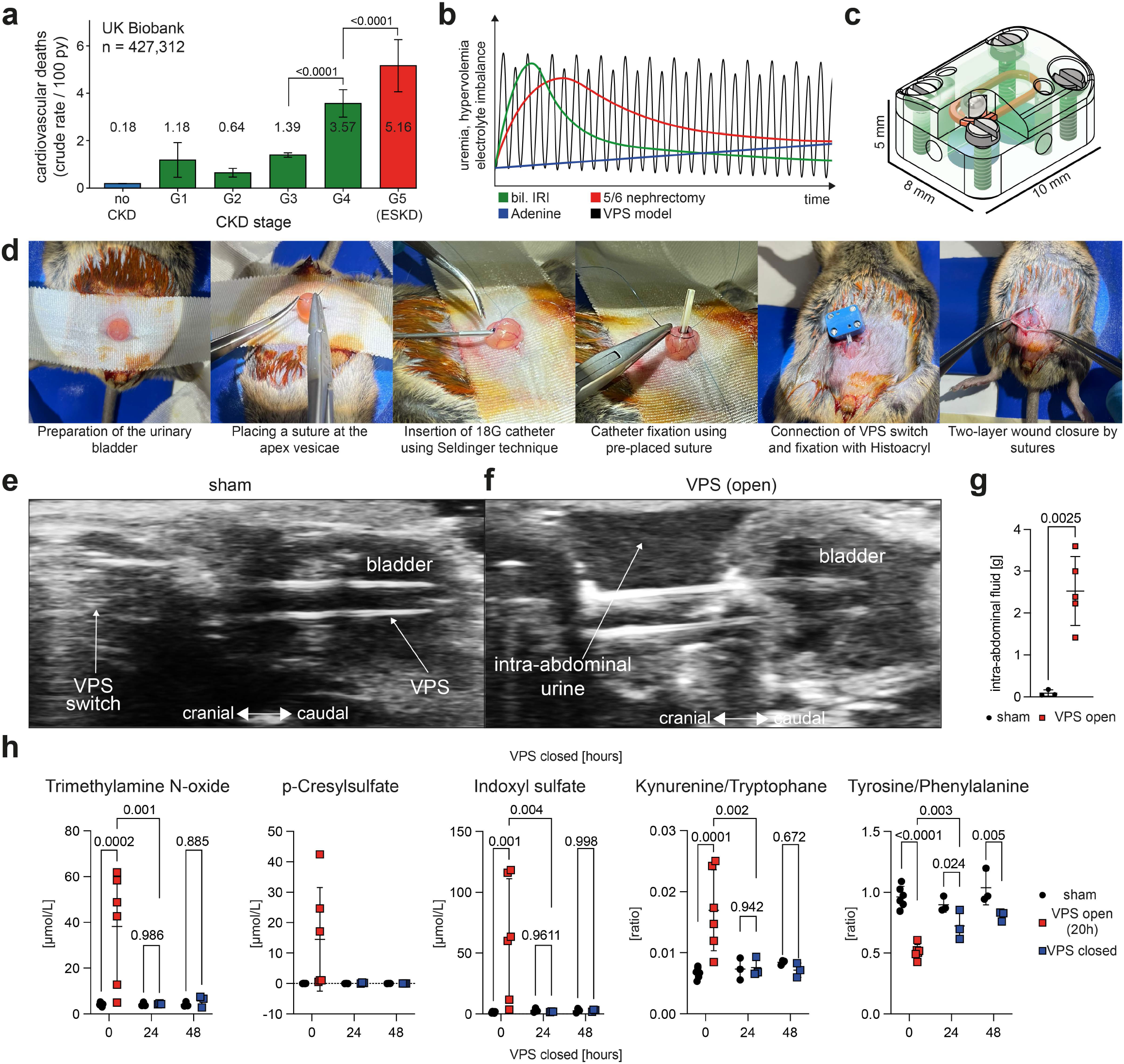
**a)**Bar chart indicating cardiovascular death rate per 100 patient years in 427,312 patients of the UK biobank stratified by CKD stage. **b)** Schematic diagram comparing temporal progression of hypervolemia, electrolyte imbalance, and uremia in current CKD mouse models and the VPS model. bil IRI: bilateral Ischemia-reperfusion injury. **c)** Technical Drawing of the VPS switch. **d)** Representative images of the surgical VPS implantation in vivo. **e-f)** Representative sonography images of the abdominal cavity with implanted VPS-switch in sham (left) and VPS (right) mice after 48h of switch opening. **g)** Quantification of the intraabdominal fluid after 48 hours of an open VPS switch comparing sham and VPS mice. **h)** Quantification of UPLC-M/MS-detected uremic toxins and amino acids in serum after 20h open shunt (0 hours VPS closed) as well as 24 and 48 hours after closing the VPS. As for p-Cresylsulfate quantification several values remained lower than the limit of quantification, no statistics are shown. *Statistics: For a univariate Cox regression. For g a two tailed t-test was performed. For h a two-way ANOVA (Tukey corrected) was calculated*

**Figure S2:**
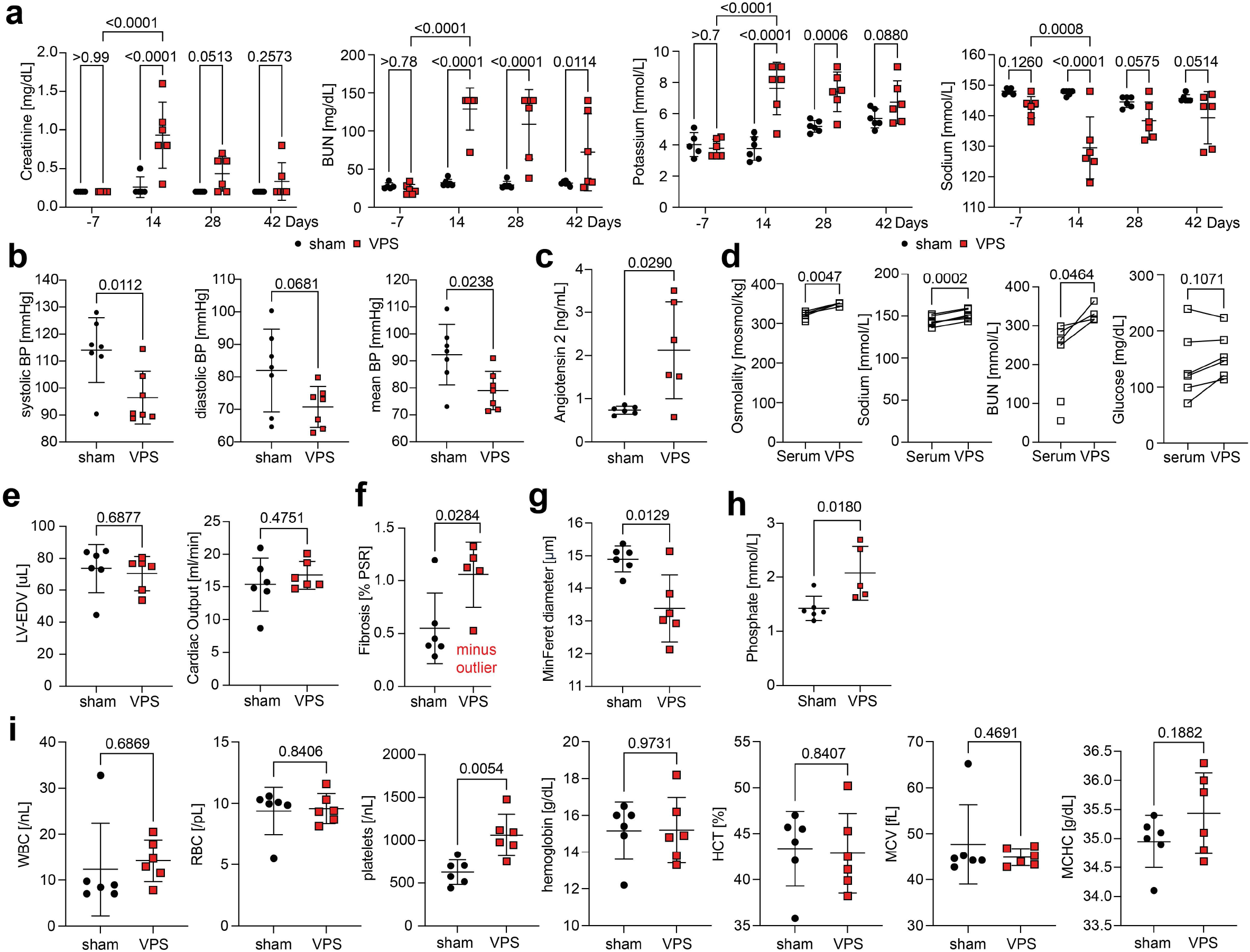
**a)**Quantification of Creatinine, BUN, potassium and sodium 7 days before and 14, 28 and 48 days after VPS surgery. **b)** Systolic, diastolic and mean arterial blood pressure in sham and VPS mice after 4 weeks of VPS. **c)** ELISA for murine angiotensin II in serum of sham and VPS mice (4 weeks VPS). **d)** Quantification of osmolality, sodium, BUN and Glucose in serum and abdominal urine of sham and VPS mice (6 weeks VPS). **e)** LV-EDV and cardiac output in sham and VPS mice (6 weeks VPS). **f)** Quantification of fibrosis as determined by PSR-positive area in hearts from sham and VPS mice without fibrotic VPS outlier. **g)** Quantification of average cardiomyocyte MinFeret in hearts from sham and VPS mice. **h)** Serum measurement of inorganic phosphate in serum samples after 6 weeks of cyclic uremia. **i)** Blood count in sham and VPS mice 4 weeks after cyclic uremia. *Statistics: For a a Two way ANOVA (Tukey corrected) was calculated. For b, c, e - g and i a Welch‘s t-test was calculated. For d a paired t-test was calculated. For h an unpaired t-test was calculated*.

**Figure S3:**
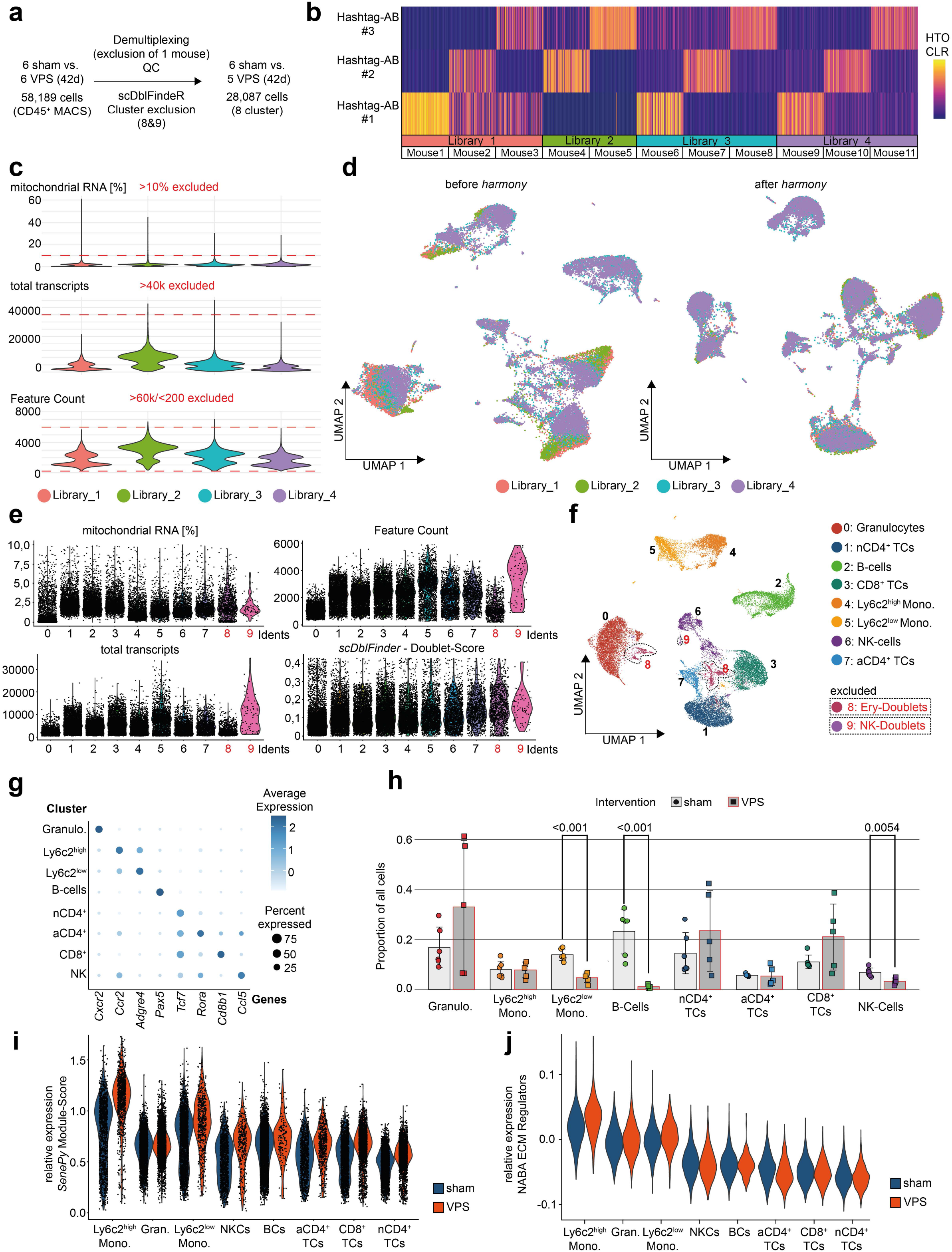
**a)**Schematic of the QC pipeline of 58,189 detected CD45^+^ immune cells (10X 3’ GEM-X) resulting in 28,087 high quality single cells after demultiplexing, quality control and doublet exclusion. **b)** Heatmap displaying the Centered Log Ratio (CLR) of the unique hashtag antibody sequence expression (hashtag sequences #1, #2, #3) in 100 random cells annotated to the individual mice over 4 multiplexed libraries. **c)** Violin Plots displaying the fraction of mitochondrial RNA, total transcripts and feature counts per cell stratified by multiplexed libraries. Cells with mitochondrial RNA content >10%, more than 40k total transcripts and less than 200 or more than 6000 features (genes) were excluded from downstream analysis. **d)** UMAP embedding before (left) and after (right) harmony integration stratified by multiplexed libraries. **e)** Violin Plots displaying the fraction of mitochondrial RNA, total transcripts, feature counts and *scDoubletfindeR*-Score per cell stratified by annotated cell clusters. **f)** UMAP embedding with annotation of the cell clustering. Clusters 8 and 9 were excluded from downstream analysis due to high expression of erythrocyte marker genes and doublet-like conformation. **g)** Dotplot displaying the top marker genes per cell cluster. **h)** Barplot displaying the proportion of each cell cluster stratified by intervention (sham vs. VPS). **i)** Violin-Plot showing relative expression of *SenePy*-defined senescence gene signatures in immune cells stratified by cluster and split by intervention. **j)** Violin-Plot showing relative expression of the *NABA Matrisome gene* signature in immune cells stratified by cluster and split by intervention. *Statistics: For h per-animal cell-type proportions were compared between sham and VPS mice using a robust empirical Bayes moderated test on arcsine-transformed proportions (propeller package)*.

