## Supplemental Figures for "Cyclic uremia without kidney injury drives cardiovascular disease with rapid onset of immunosenescence and heart failure"

Gideon JL Schaefer<sup>1,2</sup>, Patrick Droste<sup>1,2,3</sup>, Carla Schikarski<sup>1,2</sup>, Xinhui Li<sup>1,2</sup>, Marius Kohl<sup>1,2</sup>, Niklas Lutterbach<sup>4</sup>, Henriette de Loor<sup>5</sup>, Lars Koch<sup>1,2</sup>, Sylvia Menzel<sup>1,2</sup>, Ling Zhang<sup>1,2</sup>, Qingqing Long<sup>1,2</sup>, Kristian Vogt<sup>1,2</sup>, Anne-Sophie Andries<sup>1,2</sup>, Anne Babler<sup>1,2</sup>, David Schumacher<sup>2,6</sup>, KM Schneider<sup>7,8,9,12</sup>, Andreas Hoefft<sup>10</sup>, Peter Boor<sup>3</sup>, Bjorn Meijers<sup>5,11</sup>, Rebekka K Schneider<sup>4</sup>, Carolin V. Schneider<sup>12</sup>, Axel Honné<sup>13</sup>, Rafael Kramann<sup>1,2,14#</sup> & Konrad Hoefft<sup>1,2#</sup>

1) Department of Nephrology and Hypertension, Rheumatology and Immunology (Medical Clinic II), medical faculty RWTH Aachen 2) Center of Excellence in Nephrology, University Hospitals Aachen, Cologne, Dusseldorf 3) Institute of Pathology, RWTH Aachen University, Aachen, Germany 4) Department of Cell Biology, Institute for Biomedical Technologies, RWTH Aachen University, Aachen, Germany 5) Department of Microbiology, Immunology and Transplantation, Nephrology and Renal Transplantation Research Group, KU Leuven, Leuven, Belgium 6) Department of Anesthesiology, RWTH Aachen University, Aachen, Germany 7) Department of Medicine 1, University Hospital Carl Gustav Carus Dresden, TUD Dresden University of Technology, Dresden, Germany 8) Else Kroener Fresenius Center for Digital Health, Medical Faculty Carl Gustav Carus, TUD Dresden University of Technology, Dresden, Germany 9) Center for Regenerative Therapies Dresden (CRTD), TUD Dresden University of Technology, Dresden, Germany 10) Department of Anesthesiology and Intensive Care Medicine, University Hospital of Bonn, Bonn, Germany 11) Department of Nephrology and Kidney Transplantation, University Hospital Leuven, Leuven, Belgium 12) Department of Gastroenterology, Metabolic diseases and Intensive Care, University Hospital RWTH Aachen, Aachen, Germany 13) Medical Research Workshop, UK-Aachen Hospital, Germany 14) Department of Internal Medicine, Nephrology and Transplantation, Erasmus Medical Center, Rotterdam, the Netherlands, #corresponding authors

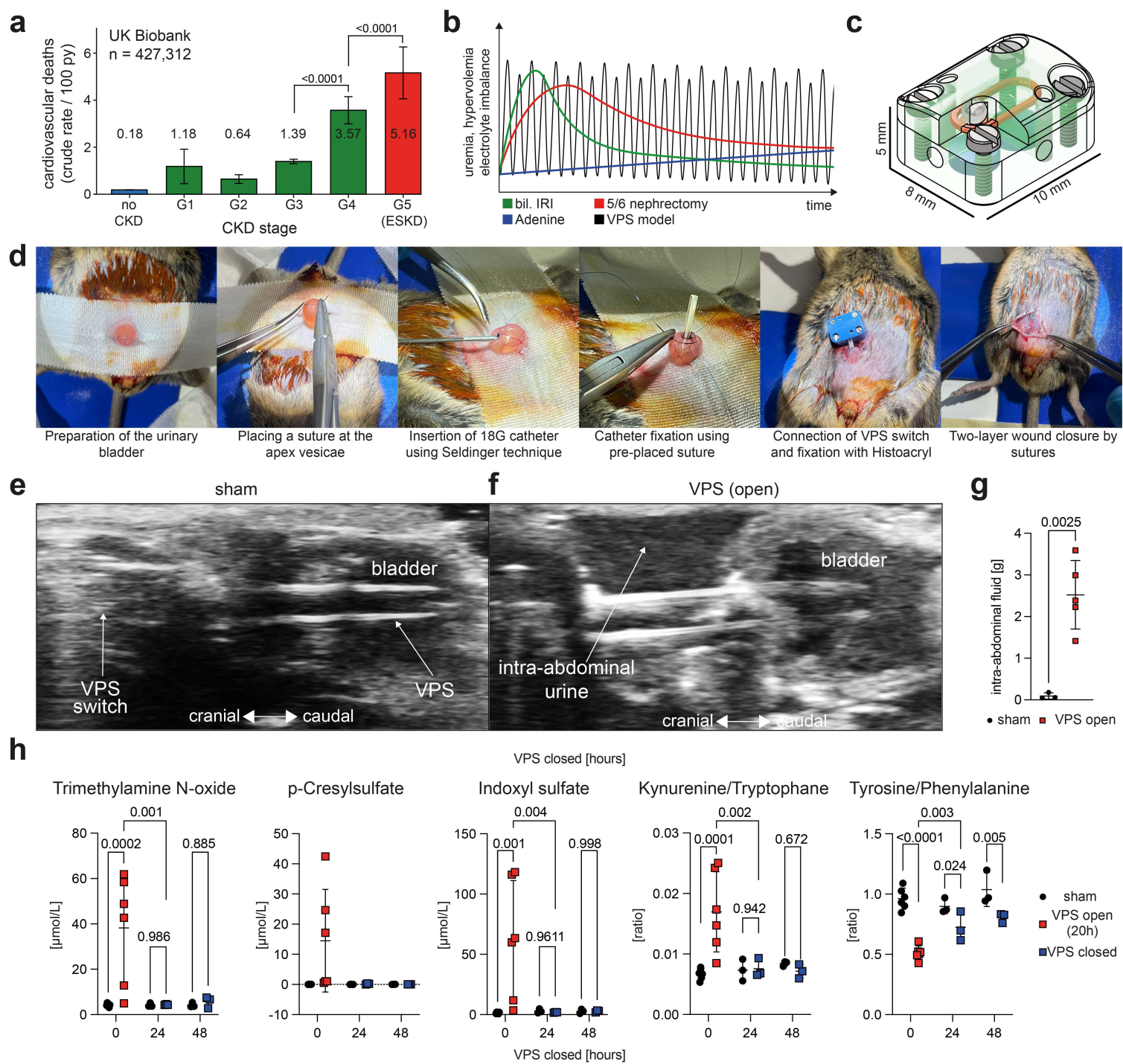

**Figure S1:** **a)** Bar chart indicating cardiovascular death rate per 100 patient years in 427,312 patients of the UK biobank stratified by CKD stage. **b)** Schematic diagram comparing temporal progression of hypervolemia, electrolyte imbalance, and uremia in current CKD mouse models and the VPS model. bil IRI: bilateral Ischemia-reperfusion injury. **c)** Technical Drawing of the VPS switch. **d)** Representative images of the surgical VPS implantation in vivo. **e-f)** Representative sonography images of the abdominal cavity with implanted VPS-switch in sham (left) and VPS (right) mice after 48h of switch opening. **g)** Quantification of the intraabdominal fluid after 48 hours of an open VPS switch comparing sham and VPS mice. **h)** Quantification of UPLC-M/MS-detected uremic toxins and amino acids in serum after 20h open shunt (0 hours VPS closed) as well as 24 and 48 hours after closing the VPS. As for p-Cresylsulfate quantification several values remained lower than the limit of quantification, no statistics are shown. *Statistics: For a univariate Cox regression. For g a two tailed t-test was performed. For h a two-way ANOVA (Tukey corrected) was calculated.*

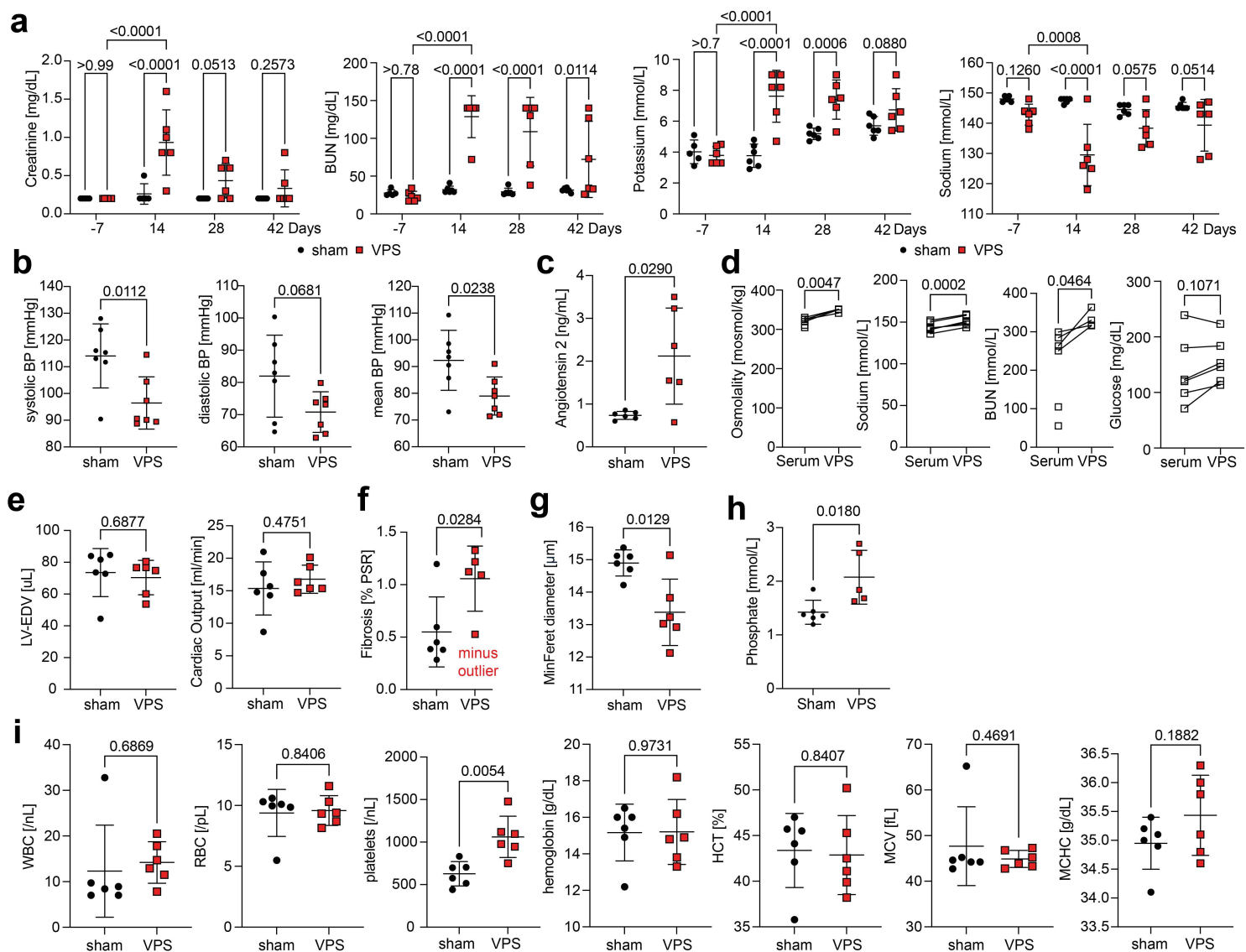

**Figure S2: a)** Quantification of Creatinine, BUN, potassium and sodium 7 days before and 14, 28 and 48 days after VPS surgery. **b)** Systolic, diastolic and mean arterial blood pressure in sham and VPS mice after 4 weeks of VPS. **c)** ELISA for murine angiotensin II in serum of sham and VPS mice (4 weeks VPS). **d)** Quantification of osmolality, sodium, BUN and Glucose in serum and abdominal urine of sham and VPS mice (6 weeks VPS). **e)** LV-EDV and cardiac output in sham and VPS mice (6 weeks VPS). **f)** Quantification of fibrosis as determined by PSR-positive area in hearts from sham and VPS mice without fibrotic VPS outlier. **g)** Quantification of average cardiomyocyte MinFeret in hearts from sham and VPS mice. **h)** Serum measurement of inorganic phosphate in serum samples after 6 weeks of cyclic uremia. **i)** Blood count in sham and VPS mice 4 weeks after cyclic uremia. *Statistics: For a a Two way ANOVA (Tukey corrected) was calculated. For b, c, e - g and i a Welch's t-test was calculated. For d a paired t-test was calculated. For h an unpaired t-test was calculated.*

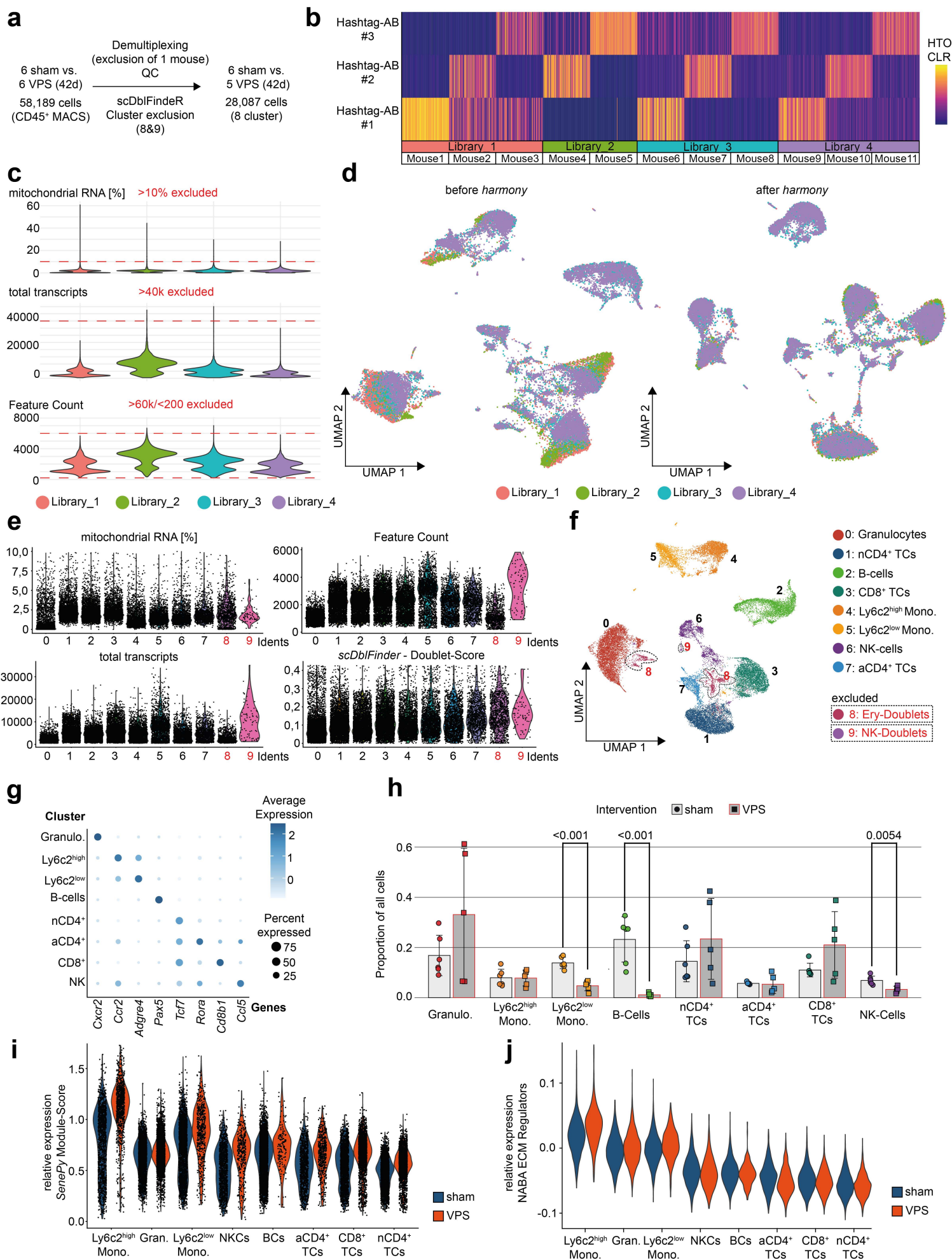

**Figure S3:** **a)** Schematic of the QC pipeline of 58,189 detected CD45<sup>+</sup> immune cells (10X 3' GEM-X) resulting in 28,087 high quality single cells after demultiplexing, quality control and doublet exclusion. **b)** Heatmap displaying the Centered Log Ratio (CLR) of the unique hashtag antibody sequence expression (hashtag sequences #1, #2, #3) in 100 random cells annotated to the individual mice over 4 multiplexed libraries. **c)** Violin Plots displaying the fraction of mitochondrial RNA, total transcripts and feature counts per cell stratified by multiplexed libraries. Cells with mitochondrial RNA content >10%, more than 40k total transcripts and less than 200 or more than 6000 features (genes) were excluded from downstream analysis. **d)** UMAP embedding before (left) and after (right) harmony integration stratified by multiplexed libraries. **e)** Violin Plots displaying the fraction of mitochondrial RNA, total transcripts, feature counts and *scDoubletfinder*-Score per cell stratified by annotated cell clusters. **f)** UMAP embedding with annotation of the cell clustering. Clusters 8 and 9 were excluded from downstream analysis due to high expression of erythrocyte marker genes and doublet-like conformation. **g)** Dotplot displaying the top marker genes per cell cluster. **h)** Barplot displaying the proportion of each cell cluster stratified by intervention (sham vs. VPS). **i)** Violin-Plot showing relative expression of *SenePy*-defined senescence gene signatures in immune cells stratified by cluster and split by intervention. **j)** Violin-Plot showing relative expression of the *NABA Matrisome* gene signature in immune cells stratified by cluster and split by intervention. *Statistics: For h per-animal cell-type proportions were compared between sham and VPS mice using a robust empirical Bayes moderated test on arcsine-transformed proportions (propeller package).*

**Supplemental Table 1:** UPLC-M/MS quantification of uremic toxins in VPS

**Supplemental Table 2:** Marker genes of *Seurat* cell clusters in peripheral immune cells

**Supplemental Table 3:** *DESeq2*-based differentially expressed genes in VPS vs. sham per *Seurat* cell cluster
